# Inter-Kingdom Interactions and Environmental Influences on the Oral Microbiome in Severe Early Childhood Caries

**DOI:** 10.1101/2024.09.26.615216

**Authors:** Lingjia Weng, Yuqi Cui, Wenting Jian, Yuwen Zhang, Liangyue Pang, Yina Cao, Yan Zhou, Wei Liu, Huancai Lin, Ye Tao

## Abstract

Dental caries arise from intricate interactions among oral microorganisms, impacting ecological stability and disease progression. This study delves into the bacterial-fungal dynamics in severe early childhood caries (S-ECC) among 61 children aged 3-4 years with complete deciduous dentition. We evaluated environmental factors such as saliva pH, buffering capacity, and trace elements (iron, fluoride). We examined the performance of 16S rRNA V1-V9 regions gene and internal transcribed spacer (ITS) primers for bacteria and fungi from plaque and saliva to characterize community compositions and diversity. Saliva pH and buffering capacity were also measured. Findings revealed significant shifts in bacterial diversity in S-ECC saliva samples, marked by decreased diversity and elevated abundance of cariogenic species, particularly *Streptococcus mutans*. *Candida albicans* was notably more prevalent in the S-ECC group, implicating its potential role in pathogenesis. Iron and fluoride concentrations showed no significant correlation with microbial community structure. Network analyses uncovered complex intra- and inter-kingdom interactions, underscoring cooperative and competitive dynamics. S-ECC children exhibited higher abundances of bacteria (*Streptococcus mutans*, *Granulicatella*, *Actinomyces*) and fungi (*Candida albicans*), with specific microbial taxa associated with reduced saliva pH.

**Importance:** This study illuminates the intricate relationship between bacteria and fungi within the oral microbial community of children, specifically highlighting differences between those with severe early childhood caries (S-ECC) and those without caries. Through an extensive analysis of the microbial composition in both saliva and dental plaque, we identified a significant increase in the abundance of specific bacterial taxa (e.g., S. mutans, Granulicatella, Actinomyces) and fungal species (e.g., C. albicans) in the oral cavities of children with S-ECC. This finding underscores the potential role of these microorganisms in the development of caries.

Contrary to previous studies that emphasize the importance of iron and fluoride in oral health, our research found no significant correlation between the concentrations of these elements and the composition of oral microbial communities. This result challenges conventional understanding and opens new avenues for future research. Additionally, we discovered an association between certain microbial species and reduced salivary pH, offering fresh insights into the relationship between the oral microenvironment and caries development.

The implications of our findings are substantial for the development of prevention and intervention strategies targeting childhood caries. They also underscore the critical need for a deeper exploration of oral microbial interactions and their environmental influences.

## Introduction

The ecological plaque hypothesis asserts that dental caries originate from a symbiotic yet dynamic consortium of microorganisms, including but not limited to fungi and bacteria, within the oral cavity (1, 2). These microorganisms contribute to oral ecological stability or disease progression via symbiotic relationships, competitive dynamics, and antagonistic interactions. Kalan et al. (2016) highlighted the intimate interactions between fungi and bacteria, which coalesce into robust three-dimensional biofilms, thereby augmenting their pathogenicity (3–5). The fungal-bacterial nexus has recently piqued interest in the pathogenetic landscape of oral infectious diseases (6–8). Contemporary research has predominantly focused on the interplay between *Candida albicans* (*C. albicans*) and various bacteria.

Notably, *Enterococcus faecalis*, frequently isolated alongside *C. albicans* in the oral cavity, has been shown to adhere to *C. albicans* hyphae, with in vitro studies demonstrating enhanced biofilm formation (9–11) and tolerance to alkaline environments, potentially exacerbating periapical inflammatory responses (12, 13). *Streptococcus mutans* (*S. mutans*), a notorious cariogenic bacterium, has been implicated in a potential synergistic relationship with *C. albicans*, particularly in the oral cavity of children with severe early childhood caries (S-ECC) (14, 15). However, reports also indicate an antagonistic relationship between the two, where compounds secreted by *S. mutans* may inhibit *C. albicans* growth(16–18). Additionally, other Streptococcus species such as *Streptococcus sanguis*, *Streptococcus gordonii*, and *Streptococcus oralis* have also been reported to synergistically enhance *C. albicans* overgrowth (19–22), although the underlying mechanisms remain to be elucidated. While numerous studies have been conducted in vitro, the intricate interplay between oral bacterial and fungal communities is not fully understood. High-throughput sequencing technologies now provide tools for dissecting these interactions at the omics level, providing a holistic view of the microecological etiology of caries.

Saliva pH, buffering capacity, and trace elements are vital for maintaining oral ecosystem stability. The salivary buffering system, exemplified by bicarbonate, ensures oral pH homeostasis. Excessive sugar intake can lead to a decrease in pH due to carbohydrate fermentation by acidogenic bacteria, resulting in tooth surface demineralization and caries development (23, 24). Research indicates that pH reduction can alter bacterial community structures and decrease diversity, with an increase in Firmicutes and Lactobacillus under acidic conditions (25, 26).

Previous studies have identified iron as a crucial nutrient for bacterial and fungal growth (27, 28), with many pathogenic bacteria possessing efficient iron uptake systems that may concurrently disadvantage beneficial bacteria (29, 30). Iron supplementation has been suggested as a means to thwart the progression of caries (31, 32). Similarly, fluoride is recognized for its role in caries prevention, primarily by hindering enamel demineralization and promoting remineralization (33–35). However, the impact of fluoride on complex in vivo biofilm communities may be less evident than previously thought.

Our previous exploratory study revealed differences in fungi from different ecological sites.(36) Here, we comprehensively analyzed the composition of the saliva and dental plaque microbiota of caries-free and S-ECC children aged between 3 and 4 years. The specific aims of our study were (i) to compare the differences in bacterial and fungal diversity between different ecological sites in the S-ECC and caries-free groups and (ii) to understand the intricate relationships within oral bacterial and fungal communities and (iii) to discover the interplay between saliva pH, iron, fluoride, and oral microbial communities.

## Materials and Methods

### 1. Population

Sixty-one children aged 3 to 4 years from two kindergartens in Zengcheng District, Guangzhou, with complete deciduous dentition, were recruited for this study. The exclusion criteria were as follows: (1) had systemic disease and had used antibiotics within 3 months; (2) had salivary gland disease and/or other oral diseases (such as periodontitis and oral mucosal disease); (3) had applied of topical and systemic fluoride within 6 months; (4) had primary teeth with enamel hypoplasia(37). Children’s general oral hygiene habits and dietary habits were investigated through structured questionnaires administered to their guardians or caregivers. Written informed consent was obtained from the guardians of all participants and ethical approval for the study was obtained from the Ethics Committee of the Hospital of Stomatology, Sun Yat-sen University, in Guangzhou, China (KQEC-2020-60-02).

### 2. Clinical Examination

The International Caries Detection and Assessment System II (ICDASII) criteria was adopted for clinical examination(36). Twenty-eight children with S-ECC [decayed, missing, or filled tooth surfaces (dmfs) ≥ 4] and 33 caries-free (dmfs = 0) children participated in the study(38). Two experienced dentists (Cui Y. and Zhang Y., kappa: 0.80-0.89) performed oral examinations using a standard mouth mirror, headlamp, and community periodontal index (CPI) probe. The visible plaque index (VPI) was used to examine all the teeth of the sample population, including the distal, medial, proximal, and lingual or palatal surfaces of each tooth under a natural light source. The results were expressed as a percentage of the total number of examined sites with visible plaque.

### 3. Sample Collection

All samples were collected between 8 a.m. and 9 a.m. (Fig. 1). Children were required to refrain from tooth brushing for 12 ± 4 h and avoid eating or drinking for 2 h before sampling.

**Fig. 1.**
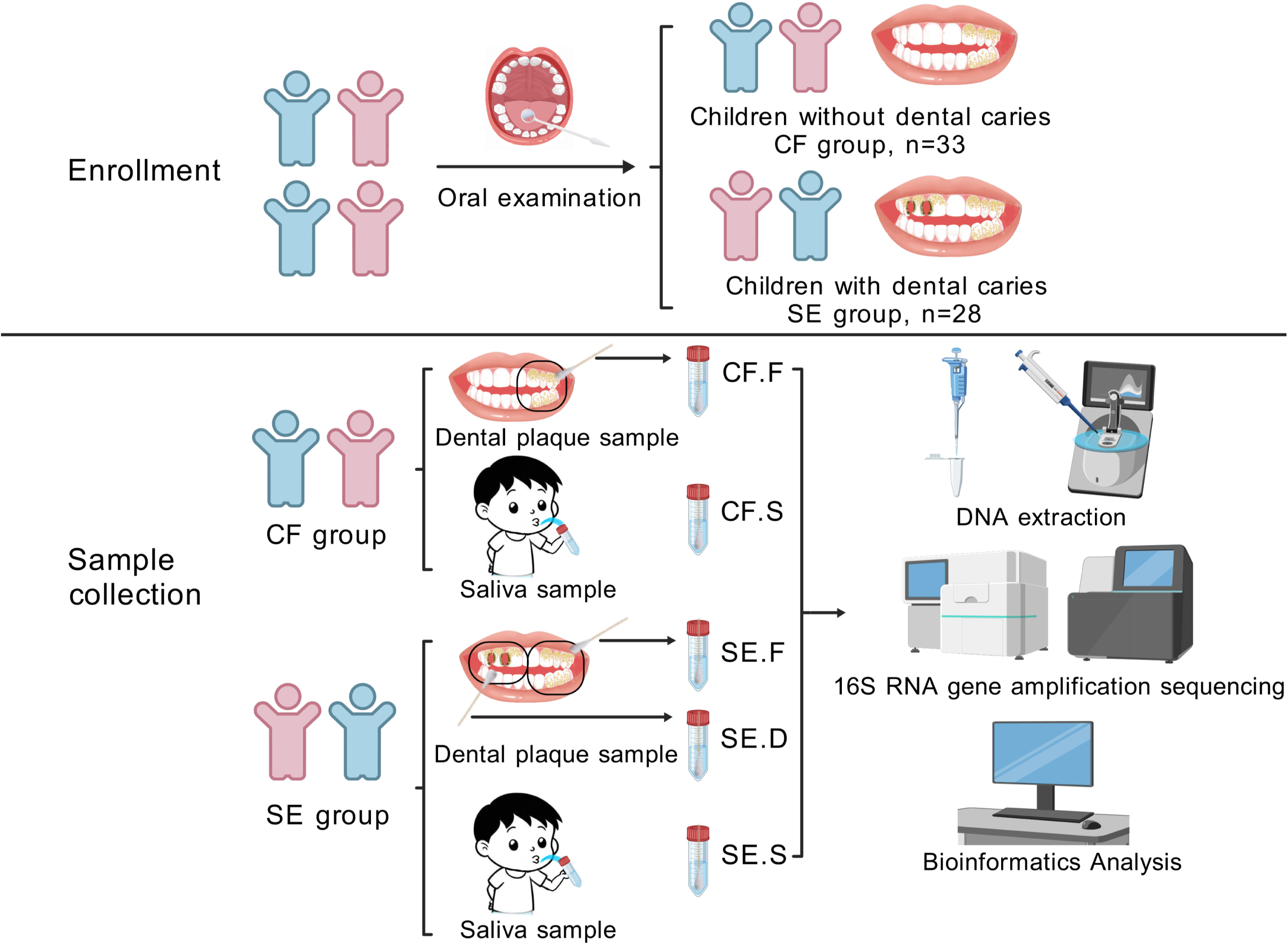
Eligible children were enrolled in the study and divided into the SE and CF groups after oral examination. In both groups, mixed supragingival plaque and nonirritating saliva were collected from available healthy tooth surfaces. In the SE group, mixed supragingival plaque from caries-damaged tooth surfaces was also collected. The samples from different groups and sections were then subjected to DNA extraction and analysis.

The pH and buffer capacity of saliva were measured as follows: a caries indicator test strip (Xierou, China) was placed in the child’s mouth, and the test strip was wetted with saliva. After the color of the test strip stabilized (usually 10 seconds), the test strip was compared with the standard colorimetric card on the packaging bottle, and the values were recorded.

Samples of supragingival plaque were collected from tooth surfaces and categorized as follows: mixed supragingival plaque from available healthy tooth surfaces in both the CF and SE groups (F), and mixed plaque from decayed tooth surfaces in the SE group (D), were collected using a sterile cotton swab. All plaque samples were then placed in DNA-free TE buffer (PH = 7.4). After all plaque samples were collected, 5ml of saliva (S) was collected using 15ml sterile centrifuge tubes from all individuals.

All biological samples were immediately placed on dry ice and transferred to a -80°C freezer for storage before further analysis. The samples collected represent five different spatial niches, three in the SE group: SE. F (n = 28), SE. D (n = 28), SE. S (n = 28), and two in the CF group: CF. F (n = 33), CF. S (n = 33).

### 4. Laboratory Methods

DNA was extracted using a DNA extraction kit (Takara Bio, China) according to the protocol of the manufacturer. The DNA concentration and purity were measured using the NanoDrop One (Thermo Fisher Scientific, USA).

For bacterial community analysis, a portion of the 16S rRNA V1-V9 regions was amplified with the barcoded universal primers (27F-AGRGTTYGATYMTGGCTCAG and (1492R-RGYTACCTTGTTACGACTT). The fungal ITS2 region was amplified using PCR. The forward primer sequence (gITS7ngs-GTGARTCATCRARTYTTTG) and the reverse primer sequence (ITS4ngs-TCCTSCGCTTATTGATATGC) were shown in our previous study. (36, 39) PCR reactions, containing 25 μl 2x Premix Taq (Takara Bio, China), 1 μl each primer(10 μM) and 3 μl DNA (20 ng/μl) template in a volume of 50 µl, were amplified by thermocycling: 5 min at 94°C for initialization; 30 cycles of 30 s denaturation at 94°C, 30 s annealing at 52°C, and 30 s extension at 72°C; followed by 10 min final elongation at 72°C. The PCR instrument was BioRad S1000 (Bio-Rad Laboratory, USA).

The length and concentration of the PCR products were detected by 1% agarose gel electrophoresis. Samples with bright main strips could be used for further experiments. PCR products were mixed in equidensity ratios according to the GeneTools Analysis Software (Version 4.03.05.0, SynGene). Then, the mixture of PCR products was purified with E.Z.N.A. Gel Extraction Kit (Omega, USA).

According to the 16S Amplification SMR Tbell® Library Preparation and NEBNext® Ultra™ II DNA Library Prep Kit for Illumina® (New England Biolabs, USA) standard process database building operations, sequencing of the constructed amplicon library using the PacBio Sequel II and Illumina Nova 6000 Nova platforms (Magigene Biotechnology Co., China)

### 5. Bioinformatics Analysis

#### 5.1. Sequencing Data Processing

The V1-V9 amplified sequences were split, corrected, format conversion, and removal of host sequences and sequences that did not match the upper primers to obtain the final valid data by using PacBio’s SMRT Link (Version 6.0). Fastp (Version 0.14.1, https://github.com/OpenGene/fastp) was used to cut the raw data from the ITS2 region amplification to obtain the valid splice fragment.

#### 5.2. OTU cluster and Species annotation

The valid data of all samples were clustered with 97% agreement using usearch software (V10, http://www.drive5.com/usearch/) and annotated with the HOMD database (http://www.homd.org). Details of the analysis of ITS2 region amplification data were shown in our former study.(36)

#### 5.3. Alpha Diversity and Beta Diversity

The alpha and beta diversity were analyzed using QIIME software (Version 1.9.1). Statistical differences in alpha and beta diversity were analyzed using Wilcoxon rank-sum tests between categories by pairwise comparison. Principle coordinates analysis (PCoA) and non-metric multi-dimensional scaling (NMDS) diagrams were drawn using R software (Version 2.15.3). Significant differences in species abundance between categories were analyzed by the LEfSe method, results were statistically significant when *P* value < 0.05.

#### 5.4. Environmental factors correlation analysis

Iron concentration and fluoride concentration were used as quantitative variables. Redundancy analysis (RDA) and Mantel test analysis were used to investigate their correlation and influence on the distribution of oral microbial communities. pH and salivary buffer capacity were categorical variables. MaAsLin (Multivariate Association with Linear Models) analysis(40, 41) was used to plot the correlation heatmap with a relative abundance threshold set at 0.01%. MaAsLin analysis is an analytical method used to effectively determine multivariate associations between clinical data and microbiome characteristics. All *P* values were corrected for multiple comparisons using FDR.

#### 5.5. Network prediction analysis

Based on the abundance of species in each group of samples, microbial species with relative abundances greater than 0.01% were selected. Then, R software was used to calculate Spearman correlation coefficient to determine correlation between species within the sample group. Bacterial and fungal species with |SpearmanCoef| > 0.6 and P values less than 0.01 were subjected to covariance network analysis.

#### 5.6. Fluorine concentration measurement

The instrument was calibrated before the assay. The gradient dilution of NaF solution was used to calibrate the fluoride ion concentration, which was calculated as y=58x-12. Saliva samples at -80°C were thawed at room temperature and then diluted for the assay.

#### 5.7. Iron concentration measurement

The standard curve of the Fe concentration measured with gradient-diluted FeNO3 solution using inductively coupled plasma mass spectrometer (ICP-MS) yielded a correlation coefficient R2 of 0.997 and the regression equation of y=752.184x+ 49.607 for Fe. The iron concentration in saliva was determined using ICP-MS.

The questionnaire data were entered into Excel and analyzed with SPSS software (Version 25.0). The data were expressed as the mean ± standard deviation. Differences were considered statistically significant when the two-sided *P* value was less than 0.05 for all statistical analyses.

## Results

### 1. Overview of the Subjects and Samples

A total of 61 children aged 3-4 years were included in this study, with a mean age of 44.4 ± 4.7 months, including 33 in the CF group and 28 in the SE group. The DMFS in the SE group ranged from 4 to 21, and the mean ±SD was 9.36±4.80. There were no significant differences in oral hygiene habits, oral hygiene status or dietary habits between the two groups. However significant differences were found in delivery mode and feeding mode within 6 months after birth (*P* < 0.05) (Table 3).

### 2. Sequencing Information

A total of 844232 valid sequences were generated by PacBio Sequel II sequencing, with an average of 6208 sequences per sample. Among all the samples, the maximum number of sequences was 12021, while the minimum number was 2603. A total of 4086821 effective sequences were generated by IlluminaNova600 sequencing, with an average of 27428 sequences per sample. Among all the samples, 74620 sequences were the most and 1271 sequences were the least.

### 3. Composition of oral bacterial and fungal communities at different spatial positions in the SE and CF groups

Bacteria: A total of 9 phyla were found in the overall bacterial community of all plaque and saliva samples: *Firmicutes, Bacteroidetes, Proteobacteria, Actinobacteria, Fusobacteria, Saccharibacteria_TM7, Spirochaetes, SR1* and *Gracilibacteria_GN02*. Among the five subgroups of the CF and SE groups, *Firmicutes* had the highest abundance (Cf. S: 34.8%, CF. F: 31.9%, SE. S: 42.9%, SE. F: 42.0%, SE. D: 40.3%). There was little difference in the abundance of each phylum among the five spatial locations (Fig. 2A).

**Fig. 2.**
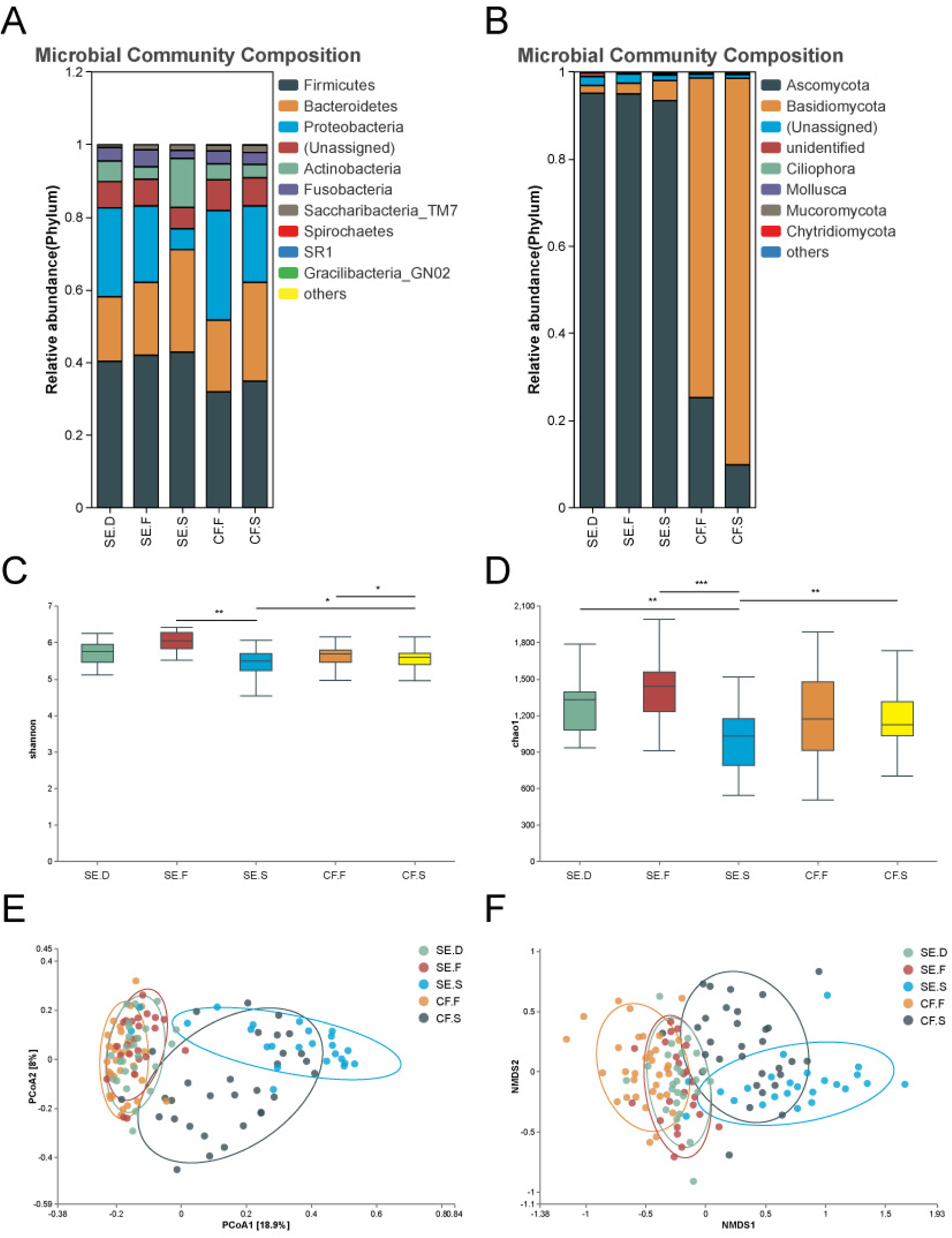
Species diversity of bacteria in various spatial locations of the oral cavity. Bacterial phylum (A) and genus (B) composition of the SE.C, SE.F, SE.S, CF.F and CF.S groups. The Alpha diversity index of each group was compared using the Shannon (C) and Chao1 indices (D). The distribution of the oral bacterial community in each group is shown through bacterial community PCoA (E) and NMDS analysis (F). The analysis revealed that the bacterial community structures of saliva and plaque samples were completely separate from each other.

A total of 15 dominant genera were identified in the overall microbial community of all plaque and saliva samples: *Streptococcus, Neisseria, Prevotella, Porphyromonas, Veillonella, Capnocytophaga, Rothia, Leptotrichia, Haemophilus, Actinomyces, Capnocytophaga, Gemella, TM7_, G-1, Fusobacterium*, and *Lautropia* (Fig. 2B).

Fungi: Two dominant phyla were found in the overall microbial community of all plaque and saliva samples: *Ascomycota* (CF.S: 9.8%, CF.F: 25.2%, SE.S: 93.4%, SE.F: 95.0%, SE.D: 95.1%) and *Basidiomycota* (CF.S: 88.7%; CF.F: 73.4%; SE.S: 4.6%, SE. F: 2.4%; SE.D: 1.8%). The abundance of the two phyla differed significantly between the SE and CF groups. The abundance of *Ascomycota* was higher in the spatial positions of the SE group, while the abundance of *Basidiomycota* was higher in the spatial positions of the CF group (Fig. 3A). Two dominant genera (relative abundance >1%) were found in the overall microbial community of all plaque and saliva samples: *Candida* (CF. S: 1.9%, CF. F: 13.7%, SE. S: 73.4%, SE. F: 73.5%, SE. D: 82.8%) and *Aspergillus* (CF. S: 0.9%, CF. F: 2.6%, SE.S: 2.2%, SE.F: 6.4%, SE.D: 1.5%) (Fig. 3B). A total of three dominant species were found in all plaque and saliva samples: *C. albicans* (CF.S: 0.3%, CF.F: 11.5%, SE.S: 59.2%, SE.F: 71.8%, SE.D: 77.1%), *Tremellomycetes sp* (CF.S: 86.2%, CF.F: 72.0%, SE.S: 0.5%, SE.F: 0.3%, SE.D. 0.2%) and *Candida tropicalis* (CF.S: 0.001%, CF.F: 0.02%, SE.F: 2.5%, SE.D: 5.2%) (Fig. 3C).

**Fig. 3.**
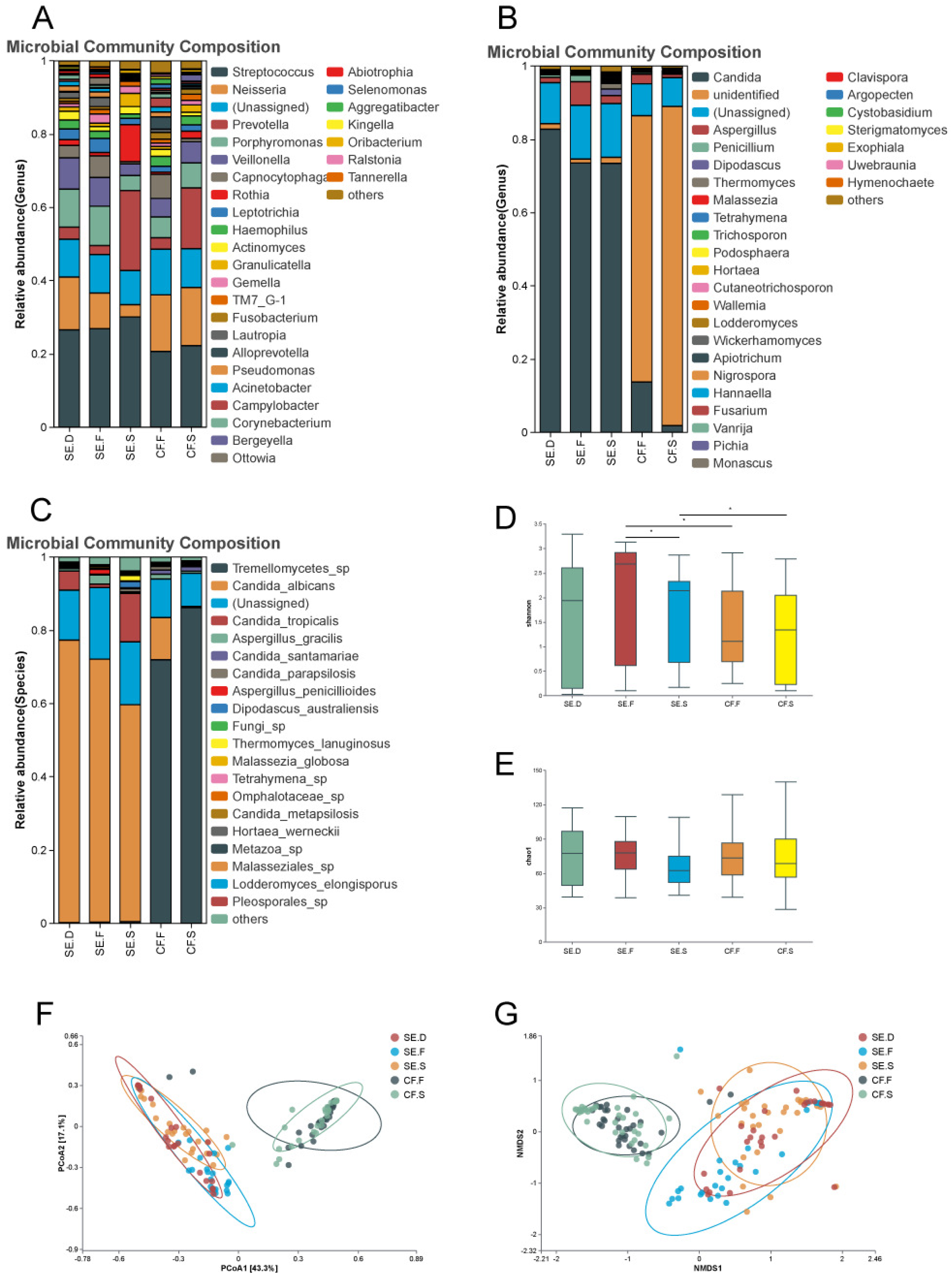
Species diversity of fungi in various spatial locations of the oral cavity. The fungal phyla (A), genera (B), and species (C) of the SE.C, SE.F, SE.S, CF.F and CF.S groups were examined. The analysis compared the Shannon index (D), and Chao1 index (E) among different groups. PCoA (F) and NMDS analysis (G) were conducted to investigate the distribution status of the oral fungal communities in each group. The results showed that saliva and plaque samples have distinct fungal community structures.

### 4. Alpha diversity analysis

Alpha (α) diversity indicators, Chao1 and Shannon were used to assess whether overall bacterial diversity differed between samples. Our statistical analyses revealed significant differences in oral bacterial community diversity among the SE.F-SE.S, SE.D-SE.S, SE.S-CF.S, and CF.F-CF.S groups (*P* < 0.05), as illustrated in Fig. 2C and D. These findings indicate a reduced bacterial diversity in saliva samples within the SE group compared to both healthy and carious plaque samples. Additionally, saliva samples from the CF group exhibited lower bacterial diversity than healthy plaque samples. Furthermore, within the SE group, bacterial diversity in saliva samples was less than that observed in the CF group.

Similarly, the Shannon and Chao1 indices revealed significant disparities in oral fungal community diversity for SE.F-SE.S, SE.F-CF.F, and SE.S-CF.S group comparisons (*P* < 0.05), as depicted in Fig. 3D and E. These results suggest that saliva samples from the SE group possessed diminished fungal diversity when compared with healthy dental plaque samples. Conversely, the fungal diversity within healthy dental plaque samples from the CF group was lower than that in the SE group. Notably, saliva samples from the SE group exhibited higher fungal diversity than those from the CF group.

### 5. Comparative analysis of the composition of oral bacterial and fungal communities at each spatial site in the SE and CF groups

Based on the NMDS and PCoA analysis using Bray-Curtis distance, we described the distribution of bacterial and fungal communities in the SE and CF groups. The results of the bacteria analysis are shown in Fig. 2E, F, and Table 1. Saliva and plaque samples were completely separated in terms of bacterial community structure. The bacterial communities in saliva samples from the SE and CF groups were significantly different, as were those in the plaque samples. Multi response permutation procedure (MRPP) statistical analysis revealed that the differences between the mentioned groups are statistically significant (*P*<0.05).

**Table 1.** MRPP analysis of differences between groups - bacteria.

| Groups | A value | *P* |
| --- | --- | --- |
| SE.D-SE.F | 0.005 | 0.066 |
| SE.D-SE.S | 0.109 | <0.001 |
| SE.F-SE.S | 0.099 | <0.001 |
| SE.F-CF.F | 0.019 | <0.001 |
| SE.S-CF.S | 0.043 | <0.001 |
| CF.F-CF.S | 0.064 | <0.001 |

The results for the fungal communities are presented in Fig. 3F, G, and Table 2. Saliva and plaque samples were completely separated in terms of fungal community structure. The fungal communities in saliva samples from the SE and CF groups were significantly different, as well as the plaque samples. MRPP statistical analysis showed that the differences between the mentioned groups were statistically significant (*P*<0.05).

**Table 2.** MRPP analysis of differences between groups - fungi.

| Groups | A value | P |
| --- | --- | --- |
| SE.D-SE.F | 0.016 | 0.036 |
| SE.D-SE.S | 0.030 | 0.008 |
| SE.F-SE.S | 0.033 | 0.005 |
| SE.F-CF.F | 0.105 | <0.001 |
| SE.S-CF.S | 0.139 | <0.001 |
| CF.F-CF.S | 0.014 | 0.061 |

The differences among the groups mentioned above are larger than the differences within each group.

### 6. Analysis of differences in oral bacterial and fungal species by spatial site in the SE and CF groups

#### 6.1. Bacteria

As shown in Fig. 4A, in the CF group, saliva samples showed a significant increase in bacteria including *Prevotella nanceiensis*, *Fusobacterium periodonticum*, *Selenomonas sp.* oral taxon 136, *Prevotella salivae*, *Neisseria sp.*, *Stomatobaculum sp* oral taxon 097, *Prevotella pallens*, and *Bergeyella sp*.

**Fig. 4.**
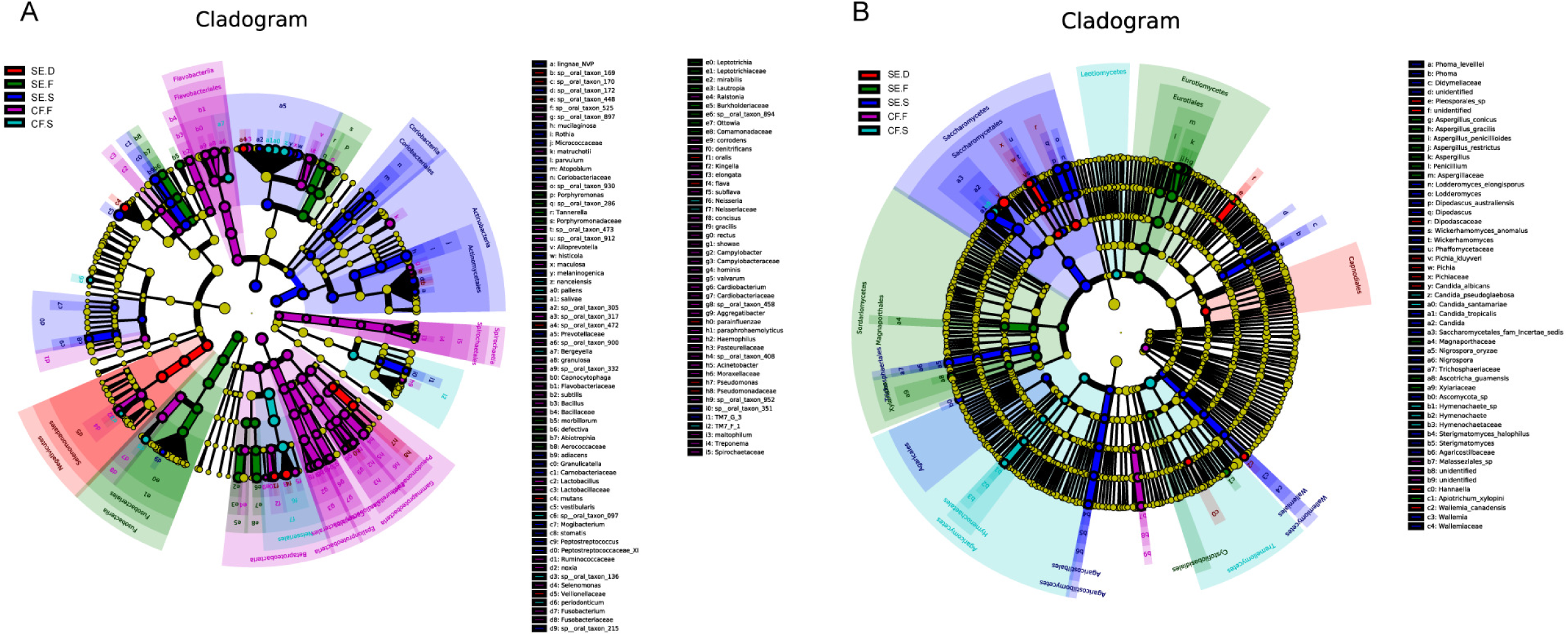
Differential bacteria (A) and fungi (B) at various sites in the oral cavities. LEfSe was used to analyze the species that differed among the groups. In the evolutionary branching diagram, circles radiating from inside to outside represent taxonomic levels from phylum to genus (or species). Each small circle at a different taxonomic level represents a taxon at that level. The size of the circle diameter is proportional to the relative abundance. Species without significant differences are uniformly colored in yellow. Different species follow the group for coloring. Different nodes are colored to represent microbial taxa that play an important role in the group. The LDA value is set to 3.

The bacteria significantly increased in healthy dental plaque samples were *Selenomonas sp*, *Acinetobacter sp*, *Porphyromonas sp* oral taxon 930, *Bacillus subtilis sp*, *Lactobacillus sp*, *Haemophilus sp*, *Kingella sp*, *Capnocytophaga granulosa sp*, *Campylobacter showae sp*, *Alloprevotella sp*, *Selenomonas noxia*, *Cardiobacterium sp*, *Corynebacterium matruchotii sp*, *Alloprevotella.sp* oral taxon 473/taxon 912, *Prevotella maculosa*, *Treponema sp*, *Actinomyces sp* oral taxon 525, *Capnocytophaga sp* oral taxon 332, *Neisseria elongata sp*, *Ralstonia sp*, *Kingella denitrificans*, *Neisseria subflava sp*, *Treponema maltophilum*, *Fusobacterium sp*, *Aggregatibacter sp*, *Aggregatibacter sp* oral taxon 458, *Campylobacter concisus*, *Cardiobacterium valvarum*, *Eikenella corrodens*, *Campylobacter rectus*, *Actinomyces sp* oral taxon 897/taxon 408, *Cardiobacterium hominis*, *Prevotella sp* oral taxon317, *Campylobacter sp*, *Bacillus sp*, *Campylobacter gracilis* and *Bergeyella sp* oral taxon 900.

In the SE group, bacteria that were significantly increased in saliva samples included *Prevotella histicola*, *Prevotella sp* oral taxon 305, *Atopobium parvulum*, *Leptotrichia sp* oral taxon 215, *Actinomyces sp* oral taxon 172, *Mogibacterium*, *Granulicatella sp*, *Atopobium sp*, *Peptostreptococcus stomatis*, *Streptococcus vestibularis sp*, *Actinomyces lingnae* NVP, *Prevotella melaninogenica*, *Rothia mucilaginosa*, *Rothia sp*, *Peptostreptococcus sp*, and *Granulicatella adiacens*. In the carious dental plaque samples, *Pseudomonas sp*, *Neisseria flava*, *S. mutans*, *Actinomyces sp* oral taxon 448/taxon 170/taxon 169, *Prevotella sp* oral taxon 448/taxon 170/taxon 169, and *S. mutans* were identified. *Prevotella sp* oral taxon 472, *Kingella oralis*, and *Haemophilus parainfluenzae* were significantly increased.

#### 6.2. Fungi

As shown in Fig. 4B, in the CF group, the significantly increased fungi in saliva samples, including *Hymenochaete sp*, *Candida pseudoglaebosa*, *Candida santamariae*, and *Hymenochaete sp*. *Malasseziales sp* were significantly increased in the healthy dental plaque samples. In the SE group, the fungi significantly increased in saliva samples, including *Nigrospora sp*, *Candida sp*, *Phoma sp*, *Lodderomyces elongisporus*, *Nigrospora oryzae*, *Phoma leveillei*, *Sterigmatomyces sp*, *Wickerhamomyces anomalus*, *Ascomycota sp*, *Sterigmatomyces halophilus*, *Dipodascus australiensis*, *Wallemia sp*, *Lodderomyces sp*, *Candida tropicalis sp*.

The fungi that were significantly more abundant in the plaque of the carious tooth surface were *Candida albicans*, *Pleosporales sp*, *Pichia kluyveri* and *Wallemia canadensis*.

### 7. Environmental factor association analysis of fungal and bacterial communities in the SE and CF groups

The values of the relevant environmental factors for the SE and CF groups are shown in Table 4.

**Table 3.** Comparison of socio-economic background and behavioral differences between the two groups.

| Variable | CF group (n=33) | SE group (n=28) |
| --- | --- | --- |
| Age (month)<sup>a</sup> | 43.55±4.80 | 45.5±4.39 |
| Weight when born<sup>b</sup> |  |  |
| < 1.5kg | 0 | 1 (3.6) |
| 1.5~2.5kg | 5 (15.2) | 3 (10.7) |
| > 2.5kg | 28 (84.8) | 24 (85.7) |
| Gestational age<sup>c</sup> |  |  |
| 28-36 weeks | 5 (15.2) | 6 (21.4) |
| > 37 weeks | 28 (84.8) | 22 (78.6) |
| Mode of delivery<sup>c</sup> |  |  |
| Cesarean section | 13 (39.4) | 21 (75.0) ** |
| Natural birth | 20 (60.6) | 7 (25.0) |
| Feeding patterns within 6 months<sup>b</sup> |  |  |
| Breastfeeding | 13 (39.4) | 15 (53.6) * |
| Mixed feeding | 18 (54.5) | 7 (25.0) |
| Pure milk powder feeding | 2 (6.1) | 6 (21.4) |
| Abstain from night feedings<sup>c</sup> |  |  |
| ≤ 2 years | 23 (69.7) | 20 (71.4) |
| > 2 years | 10 (30.3) | 8 (28.6) |
| Toothbrushing frequency<sup>c</sup> |  |  |
| ≤ 1 time/day | 23 (69.7) | 14 (50.0) |
| ≥ 2 times/day | 10 (30.3) | 14 (50.0) |
| Frequency of intake of sweets |  |  |
| 1~3 times/week | 24 (72.7) | 22 (78.6) |
| > 3 times/week | 9 (27.3) | 6 (21.4) |
| Monthly income per household<sup>c</sup> |  |  |
| < 5,000 yuan | 6 (18.2) | 6 (21.4) |
| ≥ 5,000 yuan | 27 (81.8) | 22 (78.6) |
| Visual dental plaque<sup>b</sup> |  |  |
| ≥ 20% | 31 (94.0) | 27 (96.4) |
| < 20% | 0 (0) | 1 (3.6) |
| 0% | 2 (6.0) | 0 (0) |

Values are mean ±SD or n (%)
<sup>a</sup> t-test
<sup>b</sup> Fisher's test
<sup>c</sup> Chi-square test
\* $P < 0.05$ , \*\* $P < 0.01$

**Table 4.** Environmental factors values for the CF and SE groups.

| Sample ID | Iron<br>concentration<br>(μg/L) | Fluoride<br>concentration<br>(mg/L) | Saliva pH | Buffering<br>capacity<br>(ppm) |
| --- | --- | --- | --- | --- |
| CF.1 | 2.45 | 0.060 | 6.5 | 250 |
| CF.2 | 25.26 | 0.198 | 6.5 | 250 |
| CF.3 | 10.99 | 0.090 | 5.5 | 250 |
| CF.4 | 13.88 | 0.144 | 6.0 | 250 |
| CF.5 | 10.49 | 0.102 | 7.0 | 250 |
| CF.6 | 21.87 | 0.138 | 6.0 | 250 |
| CF.7 | 15.9 | 0.072 | 6.5 | 500 |
| CF.8 | 44.96 | 0.144 | 7.0 | 500 |
| CF.9 | 41.6 | 0.096 | 6.5 | 250 |
| CF.10 | 3.98 | 0.210 | 6.0 | 250 |
| CF.11 | 15.84 | 0.096 | 6.5 | 250 |
| CF.12 | 9.44 | 0.150 | 6.5 | 250 |
| CF.13 | 10.56 | 0.060 | 5.5 | 250 |
| CF.14 | 35.24 | 0.096 | 6.5 | 500 |
| CF.15 | 15.36 | 0.132 | 6.5 | 500 |
| CF.16 | 6.25 | 0.096 | 7.0 | 500 |
| CF.17 | 14.55 | 0.162 | 6.0 | 250 |
| CF.18 | 11.29 | 0.156 | 6.0 | 250 |
| CF.19 | 19.47 | 0.174 | 7.0 | 250 |
| CF.20 | 9.67 | 0.186 | 6.5 | 250 |
| CF.21 | 25.89 | 0.174 | 6.0 | 250 |
| CF.22 | 6.85 | 0.228 | 7.0 | 500 |
| CF.23 | 11.4 | 0.096 | 7.0 | 250 |
| CF.24 | 25.28 | 0.168 | 6.0 | 250 |
| CF.25 | 12.49 | 0.144 | 6.0 | 250 |
| CF.26 | 9.74 | 0.108 | 6.5 | 250 |
| CF.27 | 10.15 | 0.186 | 6.5 | 500 |
| CF.28 | 18.27 | 0.174 | 7.0 | 500 |
| CF.29 | 10.78 | 0.108 | 7.0 | 500 |
| CF.30 | 18.66 | 0.108 | 6.5 | 250 |
| CF.31 | 7.81 | 0.120 | 8.0 | 500 |
| CF.32 | 9.4 | 0.084 | 7.0 | 500 |
| CF.33 | 5.22 | 0.150 | 6.5 | 250 |
| SE.1 | 21.41 | 0.078 | 6.5 | 500 |
| SE.2 | 32.29 | 0.072 | 6.5 | 250 |
| SE.3 | 9.66 | 0.090 | 8.0 | 500 |
| SE.4 | 15.44 | 0.090 | 7.0 | 500 |
| SE.5 | 5.17 | 0.096 | 6.5 | 250 |
| SE.6 | 34.15 | 0.084 | 5.5 | <20 |
| SE.7 | 5.78 | 0.096 | 5.5 | <20 |
| SE.8 | 27.06 | 0.150 | 7.0 | 250 |
| SE.9 | 1.95 | 0.102 | 6.5 | 500 |
| SE.10 | 11.79 | 0.132 | 5.5 | <20 |
| SE.11 | 0.65 | 0.102 | 5.5 | 500 |
| SE.12 | 9.33 | 0.132 | 5.5 | 250 |
| SE.13 | 5.12 | 0.072 | 5.5 | 500 |
| SE.14 | 27.37 | 0.066 | 6.5 | 250 |
| SE.15 | 11.76 | 0.072 | 6.0 | 250 |
| SE.16 | 13.62 | 0.108 | 6.5 | 500 |
| SE.17 | 18.02 | 0.108 | 6.5 | 500 |
| SE.18 | 57.13 | 0.084 | 5.5 | 250 |
| SE.19 | 8.91 | 0.132 | 6.0 | 250 |
| SE.20 | 22.79 | 0.078 | 6.0 | 500 |
| SE.21 | 15.41 | 0.150 | 5.5 | 250 |
| SE.22 | 11.99 | 0.084 | 6.0 | 250 |
| SE.23 | 6.46 | 0.096 | 5.5 | <20 |
| SE.24 | 14.24 | 0.084 | 5.5 | 250 |
| SE.25 | 29.74 | 0.096 | 6.0 | 250 |
| SE.26 | 17.98 | 0.096 | 8.0 | 500 |
| SE.27 | 8.5 | 0.114 | 5.5 | 250 |
| SE.28 | 14.03 | 0.126 | 5.5 | 250 |

The iron concentration, fluorine concentration, pH value and buffering capacity of the saliva of all the children were summarized.

#### 7.1. Correlation analysis of iron and fluorine levels with bacterial and fungal communities

The results of RDA (Fig. 5A, B) showed that Fe and F exhibited a negative correlation. However, there was no significant correlation between iron and fluorine levels or between the distributions of bacterial and fungal communities in the SE group and the CF group (*P* > 0.05).

**Fig. 5.**
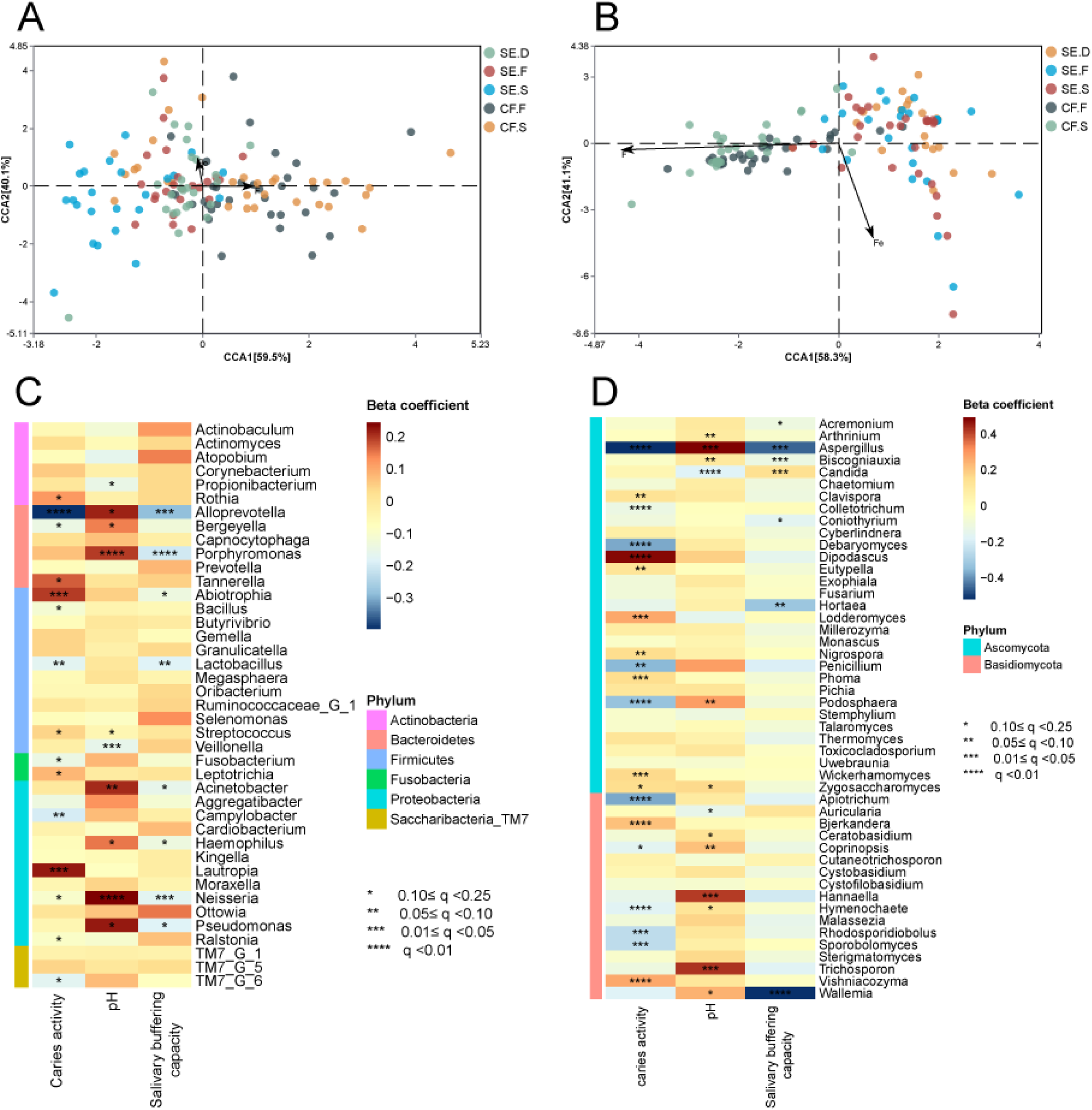
Microenvironmental factors analysis. RDA of the microbial communities and iron and fluorine concentrations at each spatial site in the SE and CF groups. (A) bacteria, (B) fungi. The arrows in the figure represent iron and fluorine, and the different dots represent the samples. The correlation is represented by the angle between the arrows. The length of the arrows represents the magnitude of the correlation, with longer arrows indicating a greater effect on the microorganisms. MaAsLin analysis of saliva pH and buffering capacity of bacterial communities (C). MaAsLin analysis of saliva pH and buffering capacity of fungal communities (D).

#### 7.2. Correlation analysis of saliva pH and buffering capacity with bacterial communities

The results of the MaAsLin analysis are shown in Fig. 5C: *Alloprevotella*, *Bergeyella sp.*, *Porphyromonas sp.*, *Acinetobacter sp.*, *Neisseria sp.* and *Pseudomonas sp.* are positively correlated with pH, while *Veillonella sp.* and *Propionibacterium sp.* are negatively correlated. *Alloprevotella sp.*, *Porphyromonas sp.* and *Lactobacillus sp.* were negatively correlated with saliva buffering capacity (*P* < 0.05).

#### 7.3. Correlation analysis of saliva pH and buffering capacity with fungal communities

The results of the MaAsLin analysis are shown in Fig. 5D: *Aspergillus sp.*, *Biscogniauxia*, *Podosphaera sp.*, *Hannaella* and *Trichosporon sp.* were positively correlated with pH, while *Candida sp* was negatively correlated with pH and positively correlated with saliva buffering capacity. The abundances of *Aspergillus sp.*, *Biscogniauxia*, *Coniothyrium sp.*, *Hortaea* and *Wallemia sp.* were significantly negatively correlated with the salivary buffering capacity(*P* < 0.05).

### 8. Network analysis and correlation analysis of bacteria and fungi

#### 8.1. Network analysis within bacteria or fungi

The analysis revealed interactions between 168 bacterial (SE.D:21, SE.F:23, SE.S:68, CF.F:22, CF.S:34) and 313 fungal genera(SE.D:71, SE.F:50, SE.S:104, CF.F:42, CF.S:46). Fig. 6A shows that the numbers of nodes and edges between bacterial communities in plaque samples were less than those in saliva samples in both the SE and CF groups. In the SE and CF groups, most bacteria-bacteria and fungi-fungi genera were positively correlated. Compared with fungi, there were more species that were negatively correlated with the bacterial communities of caries plaque. the figure shows that fungal interactions are more complex than bacterial interactions.

**Fig. 6.**
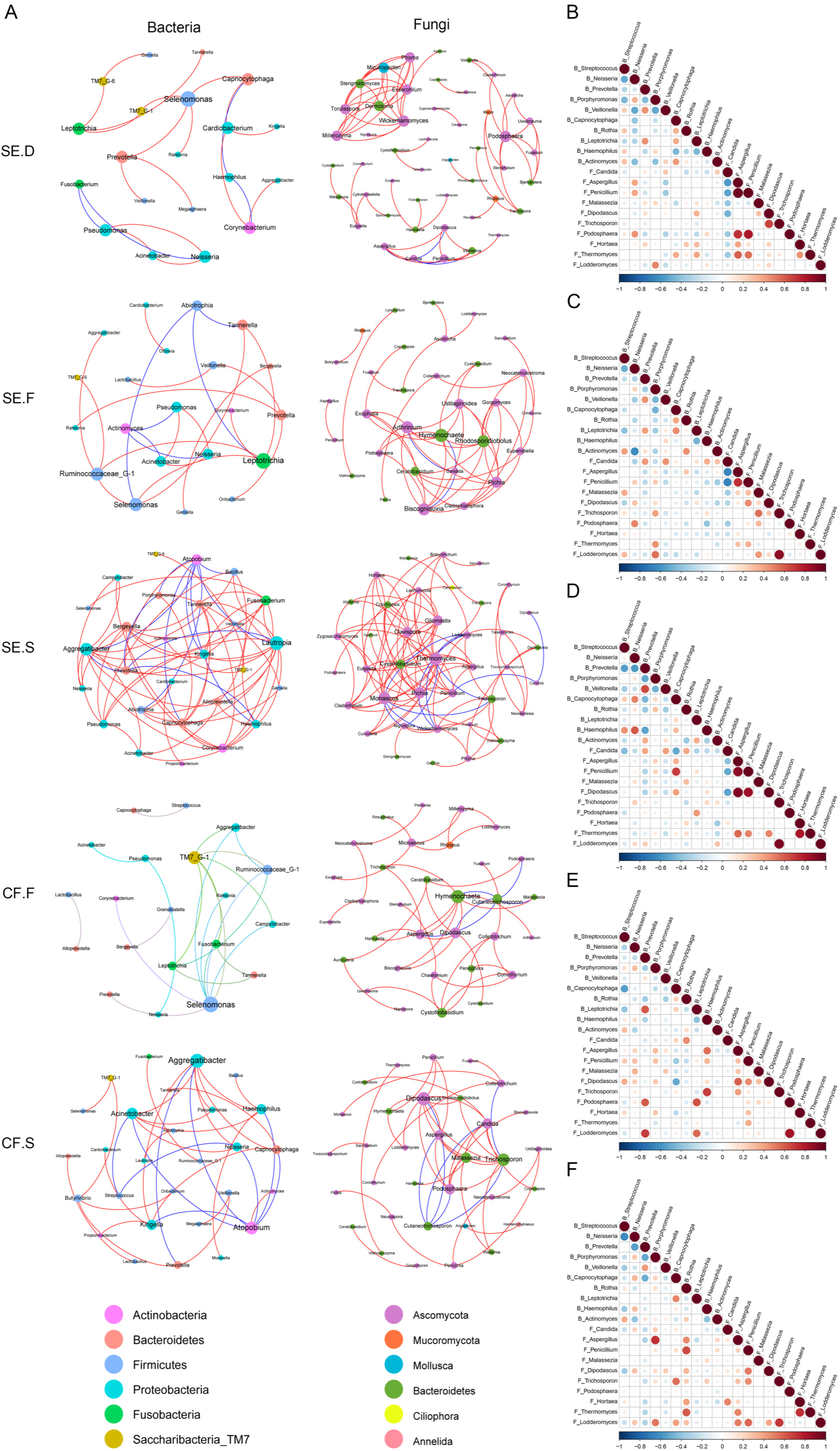
Network analysis and correlation analysis of bacteria and fungi. Network analysis of bacteria and fungi within each site of the SE group and CF group (A). A line between two species indicates a significant correlation (Spearman’s rank correlation *ρ*>0.6, *P*<0.01). The greater the number of lines between nodes and the larger the radius of the nodes are, the greater the correlation. Red represents a positive correlation between species, and blue represents a negative correlation between species. Correlation analysis of bacteria and fungi within each site of the SE group and CF group (B. SE.D, C. SE.F, D. SE.S, E. CF.F, F. CF.S). “B_ stands for bacteria” and “F_ stands for fungi”. Red and blue represent positive and negative correlations, respectively. The color shade and size of the circles are proportional to the correlation coefficient, and white represents no correlation.

#### 8.2. Correlation analysis of bacteria and fungi

Correlation analysis investigated the relationships between important bacteria and fungi at each spatial location in the SE and CF groups. Fig. 6B-F show the correlations among the five spatial sites. The results revealed significant differences in the correlation patterns between bacterial and fungal species within each spatial location. For instance, in group SE.D, *Streptococcus sp.* exhibited negative correlations with *Neisseria sp., Prevotella sp.*, *Porphyromonas sp*., *Malasseziales sp., Candida sp., and Hortaea,* but showed positive correlations with *Veillonella sp., Rothia sp., Actinomyces sp., Aspergillus sp., Penicillium sp., and Trichosporon sp*. However, in groups SE.F and SE.S, *Streptococcus sp.* positively correlated with *Malassezia sp.* and *Hortaea*.

## Discussion

### Overall trends of flora

Our research revealed a significant difference in the bacterial and fungal communities in the SE group, which included samples of healthy dental plaque, dental caries plaque, and saliva, compared to those in the CF group, which consisted of healthy dental plaque and saliva samples. Interestingly, within the CF group, no significant difference was observed in the fungal compositions of healthy dental plaque and saliva samples (*P* > 0.05). However, we found that the SE group showed distinct bacterial and fungal community structures in healthy and carious dental plaque compared to those in saliva, indicating that the occurrence of carious lesions causes changes in the microbial community configurations at each respective site.

### Dominant flora in different caries states

*C. albicans* exhibited the highest fungal abundance in the SE group, with percentages reaching 59.2% in SE.S, 71.8% in SE.F, and 77.1% in SE.D. This finding aligns with previous studies (36) that highlighted *C. albicans* as one of the most frequently cultured oral fungi, which has unique potential for causing dental caries. *C. albicans* can invade dentin tubules, leading to the demineralization and dissolution of teeth (42, 43). A meta-analysis conducted by Xiao et al. (44) based on nine cross-sectional epidemiological studies revealed that children who tested positive for *C. albicans* had a five-fold higher risk of early childhood caries (ECC) than those who tested negative. Furthermore, *C. albicans* can collaborate with other bacteria and fungi, enhancing biofilm formation, aggravating periapical inflammation when coexisting with *Enterococcus faecalis* (11–13), and amplifying its cariogenic capability when present alongside *S. mutans*. (14, 15, 45–47).

In the CF group, *Tremellomycetes sp*. was found to be the most abundant fungus, accounting for 86.2% in CF.S and 72.0% in CF.S. Notably, this fungus has not been previously reported in any studies. Therefore, more research is needed to determine whether this species is transient or permanent within the oral cavity. Furthermore, *Malasseziales sp.* was significantly higher in healthy dental plaque samples (*P* < 0.05). Previous studies have indicated that *Malasseziales sp.* is an important oral symbiotic bacterium present in both saliva and healthy dental plaque (48–51). These findings suggest that it may play a significant role in maintaining oral microecological health. Moreover, studies have shown that a diet low in carbohydrates and high in lipids, along with good oral hygiene practices, favors the growth of *Malassezia* (52).

In terms of bacterial composition, the abundance of *S. mutans* significantly increased in the saliva of the SE group (*P* < 0.05). Furthermore, notable increases were observed in bacteria belonging to *Granulicatella* and *Actinomyces* (oral taxon 448, taxon 170, and taxon 169) within the SE group. *Granulicatella* is a facultative anaerobic Gram-positive cocci bacteria (53) that can ferment glucose to produce lactic acid. Previous studies have established a link between *Granulicatella* and caries, particularly S-ECC (54, 55). Nevertheless, the intricate mechanism underlying caries formation involving *Granulicatella* remains elusive. On the other hand, *Actinomyces* is recognized as a significant pathogenic microorganism that contributes to root surface caries. *Actinomyces viscosus* has a swift ability to adhere to the root surface and bind to the collagen fibers of dentin and cementum, playing a pivotal role in regulating mineral loss on the tooth surface (56–58).

The intricate relationships within oral bacterial and fungal communities. The correlation analysis of bacteria and fungi revealed complex interactions between these two microbial groups in the SE and CF groups. Notably, fungal communities exhibited a more complex network of interactions than did bacterial communities. These results suggest that bacteria and fungi in the oral microenvironment engage in intricate interactions that contribute to the stability of the oral ecosystem or participate in pathogenic processes, which aligns with previous research findings (1, 2, 59, 60). Our study further indicated that the relationship between these two groups may reverse during the transition from a healthy to a carious state. For instance, *Streptococcus* was negatively correlated with *Hortaea* in carious plaque samples but positively correlated in healthy plaque and saliva samples. This intriguing discovery requires further identification and verification to elucidate its underlying mechanisms.

### Microenvironmental factors and oral microbial communities pH, Saliva iron and fluoride concentrations

Previous studies have shown a negative correlation between salivary pH and the decayed, missing, filled teeth (DMFT) index (61). The dental plaque pH and salivary pH are important environmental factors that affect the microenvironment of dental biofilms (25). The development of caries is associated with a decrease in pH in the microenvironment and an increase in acid-producing and acid-resistant bacteria (62, 63).

In our study, we simultaneously examined the salivary pH, which reflects the acid-producing ability of dental plaque. We found a negative correlation between the pH value and the abundances of *Veillonella, Propionibacterium*, and *Candida*. *Veillonella* is among the earliest colonizers of dental plaque and utilizes lactic acid produced by other bacteria, such as *S. mutans*, as a carbon source and produces substances such as acetic acid (64, 65). Studies have demonstrated that genes involved in lactic acid and succinic acid catabolism, as well as histidine biosynthesis, are upregulated in caries tissue samples compared to healthy saliva samples. This upregulation may contribute to the survival of *Veillonella* in an acidic environment (66).

*Propionibacterium* has been reported to be a dominant factor in dentin caries (67, 68). Studies have highlighted the significant role of *Propionibacterium* in caries progression (69). However, Chhour et al.(70) used similar molecular techniques to determine the microbial diversity in adults with advanced caries and found that an abundance of *Propionibacterium* species was not commonly detected (70). Therefore, the relationship between *Propionibacterium* and caries requires further investigation.

Recent research on *Candida* has demonstrated its status as an acid-producing and acid-tolerant bacterium, capable of independently causing caries (71–73). *Candida*-dominated communities have been observed more frequently in individuals who smoke, use removable dental prostheses, suffer from active caries, exhibit higher plaque index scores, or take steroids (52). Microscopic in situ evaluations of intact clinical biofilm samples from caries-affected individuals have suggested a pivotal role for *Candida* in the biofilm development of cariogenic bacteria. These studies revealed fungal-bacterial interactions, with images showing *streptococci* arranging around *C. albicans* hyphae, and forming distinct fungal-bacterial corncobs. (45, 74–76). Furthermore, our study revealed a negative correlation between iron and fluoride concentrations. However, there was no significant difference in the distribution of oral bacterial and fungal communities associated with iron and fluoride concentrations in either group. Previous studies have identified iron as an essential nutrient for bacteria and fungi (27, 28). Animal experiments have suggested that iron may inhibit the progression of dental caries (31, 32). Clinical studies have demonstrated an association between low iron concentration and high DMFT (61). However, the underlying mechanisms have not been fully elucidated.

Fluoride is a highly effective element for preventing early childhood caries (ECC) (33). It works by inhibiting enamel demineralization and promoting remineralization, which helps prevent cavities (34, 35). The antimicrobial and anticaries properties of fluoride are mainly due to its ability to reduce the acid tolerance of glycolysis in cariogenic bacteria residing within dental plaque (77). Nevertheless, similar to our study, fluoride coating did not have a significant effect on oral organisms in preschool children (78). A possible explanation is that while many organisms are sensitive to fluoride in free or simple biofilm models in vitro, their responses may vary within complex oral biofilm communities (77, 79).

### Conclusions

To date, limited research has been conducted on the relationships between salivary pH, iron, fluorine, and oral microbial communities. More studies are necessary to investigate the connections between these factors and oral bacterial and fungal communities.

Our research discovered differences in the structures of fungal and bacterial communities present in the oral saliva and dental plaque of children suffering from severe early childhood caries (S-ECC). The distribution of these communities varied depending on the state of caries. We identified complex interactions among fungi and bacteria present in both healthy and carious states. S-ECC children had significantly higher levels of bacteria such as *S. mutans*, *Granulicatella*, and *Actinomyces*, as well as fungi such as *C. albicans* than caries-free children. We found no significant correlation between the concentrations of iron and fluoride and the oral bacterial and fungal communities (*P* > 0.05).

Bacteria such as *Veillonella, Propionibacterium,* and *Leptotrichia* as well as fungi including *Nigrospora, Candida*, and *Bjerkandera*, were found to be associated with a decrease in saliva pH value.

## Abbreviations

C. albicans: Candida albicans
S. mutans: Streptococcus mutans
S-ECC: severe early childhood caries
CF: caries free
DMFS: decayed, missing and filled tooth surface
ICDASII: International Caries Detection and Assessment System II
SE.D: mixed plaque from decayed teeth in the severe early childhood caries group
SE.F: healthy supragingival mixed plaque in the severe early childhood caries group
SE.S: saliva samples in the severe early childhood caries group
CF.F: healthy supragingival mixed plaque in the caries-free group
CF.S: saliva samples in the caries-free group
OUT: operational taxonomic units
ITS: internal transcribed spacer
PCoA: principal co-ordinates analysis
NMDS: non-metric multi-dimensional scaling
MRPP: multi response permutation procedure
PCR: polymerase chain reaction
DNA: deoxyribonucleic acid
TE: tris-EDTA buffer
LEfSe: linear discriminant analysis effect size

## Data availability

The datasets generated during the current study are available in the SRA repository with the accession numbers PRJNA1080488 and PRJNA848441.

## Acknowledgments

The authors express gratitude to all participants of the study.

## Authors’ contributions

WLJ, CYQ and JWT contributed equally to the study. WLJ: data analysis, manuscript preparation. CYQ: oral examination, data analysis. JWT: laboratory experiment, data analysis. ZYW: oral examination, sample processing. PLY, CYN and ZY: recruitment and consent, sample and data collection. LW: data analysis, figures and tables production. LHC and TY: study design and supervision, manuscript preparation. All authors read and approved the final manuscript.

## Declarations

### Ethics approval and consent to participate

This research protocol has been reviewed and approved by the Ethics Committee of the Affiliated Stomatological Hospital of Sun Yat-Sen University (KQEC-2020-60-02). Signed informed consent was obtained from all participants; samples were processed anonymously.

### Consent for publication

All the participants in the present study consented independently. All the personal data generated during the research were kept confidential.

## References

1. Anderson AC, Rothballer M, Altenburger MJ, Woelber JP, Karygianni L, Lagkouvardos I, Hellwig E, Al-Ahmad A. 2018. In-vivo shift of the microbiota in oral biofilm in response to frequent sucrose consumption. Sci Rep 8:14202.

2. Rosier BT, De Jager M, Zaura E, Krom BP. 2014. Historical and contemporary hypotheses on the development of oral diseases: are we there yet? Front Cell Infect Microbiol 4:92.

3. Kalan L, Loesche M, Hodkinson BP, Heilmann K, Ruthel G, Gardner SE, Grice EA. 2016. Redefining the Chronic-Wound Microbiome: Fungal Communities Are Prevalent, Dynamic, and Associated with Delayed Healing. mBio 7.

4. Hoarau G, Mukherjee PK, Gower-Rousseau C, Hager C, Chandra J, Retuerto MA, Neut C, Vermeire S, Clemente J, Colombel JF, Fujioka H, Poulain D, Sendid B, Ghannoum MA. 2016. Bacteriome and Mycobiome Interactions Underscore Microbial Dysbiosis in Familial Crohn’s Disease. mBio 7.

5. Costerton JW, Stewart PS, Greenberg EP. 1999. Bacterial biofilms: a common cause of persistent infections. Science 284:1318–22.

6. Du Q, Ren B, He J, Peng X, Guo Q, Zheng L, Li J, Dai H, Chen V, Zhang L, Zhou X, Xu X. 2021. Candida albicans promotes tooth decay by inducing oral microbial dysbiosis. Isme j 15:894–908.

7. Janus MM, Crielaard W, Volgenant CM, van der Veen MH, Brandt BW, Krom BP. 2017. Candida albicans alters the bacterial microbiome of early in vitro oral biofilms. J Oral Microbiol 9:1270613.

8. Xiao J, Grier A, Faustoferri RC, Alzoubi S, Gill AL, Feng C, Liu Y, Quivey RG, Kopycka-Kedzierawski DT, Koo H, Gill SR. 2018. Association between Oral Candida and Bacteriome in Children with Severe ECC. J Dent Res 97:1468–1476.

9. Kovac J, Kovac D, Slobodnikova L, Kotulova D. 2013. Enterococcus faecalis and Candida albicans in the dental root canal and periapical infections. Bratisl Lek Listy 114:716–20.

10. Dahlén G, Blomqvist S, Almståhl A, Carlén A. 2012. Virulence factors and antibiotic susceptibility in enterococci isolated from oral mucosal and deep infections. J Oral Microbiol 4.

11. Siqueira JF, Jr., Rôças IN. 2009. Diversity of endodontic microbiota revisited. J Dent Res 88:969–81.

12. Fox EP, Cowley ES, Nobile CJ, Hartooni N, Newman DK, Johnson AD. 2014. Anaerobic bacteria grow within Candida albicans biofilms and induce biofilm formation in suspension cultures. Curr Biol 24:2411–6.

13. Du Q, Yuan S, Zhao S, Fu D, Chen Y, Zhou Y, Cao Y, Gao Y, Xu X, Zhou X, He J. 2021. Coexistence of Candida albicans and Enterococcus faecalis increases biofilm virulence and periapical lesions in rats. Biofouling 37:964–974.

14. Thomas A, Mhambrey S, Chokshi K, Chokshi A, Jana S, Thakur S, Jose D, Bajpai G. 2016. Association of Oral Candida albicans with Severe Early Childhood Caries - A Pilot Study. J Clin Diagn Res 10:Zc109–12.

15. Koo H, Bowen WH. 2014. Candida albicans and Streptococcus mutans: a potential synergistic alliance to cause virulent tooth decay in children. Future Microbiol 9:1295–7.

16. Khan F, Bamunuarachchi NI, Pham DTN, Tabassum N, Khan MSA, Kim YM. 2021. Mixed biofilms of pathogenic Candida-bacteria: regulation mechanisms and treatment strategies. Crit Rev Microbiol 47:699–727.

17. Jarosz LM, Deng DM, van der Mei HC, Crielaard W, Krom BP. 2009. Streptococcus mutans competence-stimulating peptide inhibits Candida albicans hypha formation. Eukaryot Cell 8:1658–64.

18. Vílchez R, Lemme A, Ballhausen B, Thiel V, Schulz S, Jansen R, Sztajer H, Wagner-Döbler I. 2010. Streptococcus mutans inhibits Candida albicans hyphal formation by the fatty acid signaling molecule trans-2-decenoic acid (SDSF). Chembiochem 11:1552–62.

19. Förster TM, Mogavero S, Dräger A, Graf K, Polke M, Jacobsen ID, Hube B. 2016. Enemies and brothers in arms: Candida albicans and gram-positive bacteria. Cell Microbiol 18:1709–1715.

20. Montelongo-Jauregui D, Saville SP, Lopez-Ribot JL. 2019. Contributions of Candida albicans Dimorphism, Adhesive Interactions, and Extracellular Matrix to the Formation of Dual-Species Biofilms with Streptococcus gordonii. mBio 10.

21. Jenkinson HF, Lala HC, Shepherd MG. 1990. Coaggregation of Streptococcus sanguis and other streptococci with Candida albicans. Infect Immun 58:1429–36.

22. Xu H, Sobue T, Bertolini M, Thompson A, Dongari-Bagtzoglou A. 2016. Streptococcus oralis and Candida albicans Synergistically Activate μ-Calpain to Degrade E-cadherin From Oral Epithelial Junctions. J Infect Dis 214:925–34.

23. Pyati SA, Naveen Kumar R, Kumar V, Praveen Kumar NH, Parveen Reddy KM. 2018. Salivary Flow Rate, pH, Buffering Capacity, Total Protein, Oxidative Stress and Antioxidant Capacity in Children with and without Dental Caries. J Clin Pediatr Dent 42:445–449.

24. Larsen MJ, Jensen AF, Madsen DM, Pearce EI. 1999. Individual variations of pH, buffer capacity, and concentrations of calcium and phosphate in unstimulated whole saliva. Arch Oral Biol 44:111–7.

25. Takahashi N, Nyvad B. 2011. The role of bacteria in the caries process: ecological perspectives. J Dent Res 90:294–303.

26. Kianoush N, Adler CJ, Nguyen KA, Browne GV, Simonian M, Hunter N. 2014. Bacterial profile of dentine caries and the impact of pH on bacterial population diversity. PLoS One 9:e92940.

27. Wooldridge KG, Williams PH. 1993. Iron uptake mechanisms of pathogenic bacteria. FEMS Microbiol Rev 12:325–48.

28. Tripathi A, Liverani E, Tsygankov AY, Puri S. 2020. Iron alters the cell wall composition and intracellular lactate to affect Candida albicans susceptibility to antifungals and host immune response. J Biol Chem 295:10032–10044.

29. Simonyté Sjödin K, Domellöf M, Lagerqvist C, Hernell O, Lönnerdal B, Szymlek-Gay EA, Sjödin A, West CE, Lind T. 2019. Administration of ferrous sulfate drops has significant effects on the gut microbiota of iron-sufficient infants: a randomised controlled study. Gut 68:2095–2097.

30. Jaeggi T, Kortman GA, Moretti D, Chassard C, Holding P, Dostal A, Boekhorst J, Timmerman HM, Swinkels DW, Tjalsma H, Njenga J, Mwangi A, Kvalsvig J, Lacroix C, Zimmermann MB. 2015. Iron fortification adversely affects the gut microbiome, increases pathogen abundance and induces intestinal inflammation in Kenyan infants. Gut 64:731–42.

31. Eshghi A, Kowsari-Isfahan R, Rezaiefar M, Razavi M, Zeighami S. 2012. Effect of iron containing supplements on rats’ dental caries progression. J Dent (Tehran) 9:14–9.

32. Pecharki GD, Cury JA, Paes Leme AF, Tabchoury CP, Del Bel Cury AA, Rosalen PL, Bowen WH. 2005. Effect of sucrose containing iron (II) on dental biofilm and enamel demineralization in situ. Caries Res 39:123–9.

33. Twetman S, Dhar V. 2015. Evidence of Effectiveness of Current Therapies to Prevent and Treat Early Childhood Caries. Pediatr Dent 37:246–53.

34. Bossù M, Saccucci M, Salucci A, Di Giorgio G, Bruni E, Uccelletti D, Sarto MS, Familiari G, Relucenti M, Polimeni A. 2019. Enamel remineralization and repair results of Biomimetic Hydroxyapatite toothpaste on deciduous teeth: an effective option to fluoride toothpaste. J Nanobiotechnology 17:17.

35. Walsh T, Worthington HV, Glenny AM, Marinho VC, Jeroncic A. 2019. Fluoride toothpastes of different concentrations for preventing dental caries. Cochrane Database Syst Rev 3:Cd007868.

36. Cui Y, Wang Y, Zhang Y, Pang L, Zhou Y, Lin H, Tao Y. 2021. Oral Mycobiome Differences in Various Spatial Niches With and Without Severe Early Childhood Caries. Frontiers in Pediatrics 9.

37. Blostein F, Bhaumik D, Davis E, Salzman E, Shedden K, Duhaime M, Bakulski KM, McNeil DW, Marazita ML, Foxman B. 2022. Evaluating the ecological hypothesis: early life salivary microbiome assembly predicts dental caries in a longitudinal case-control study. Microbiome 10:240.

38. Drury TF, Horowitz AM, Ismail AI, Maertens MP, Rozier RG, Selwitz RH. 1999. Diagnosing and reporting early childhood caries for research purposes. A report of a workshop sponsored by the National Institute of Dental and Craniofacial Research, the Health Resources and Services Administration, and the Health Care Financing Administration. J Public Health Dent 59:192–7.

39. Nilsson RH, Anslan S, Bahram M, Wurzbacher C, Baldrian P, Tedersoo L. 2019. Mycobiome diversity: high-throughput sequencing and identification of fungi. Nat Rev Microbiol 17:95–109.

40. Thingholm LB, Rühlemann MC, Koch M, Fuqua B, Laucke G, Boehm R, Bang C, Franzosa EA, Hübenthal M, Rahnavard A, Frost F, Lloyd-Price J, Schirmer M, Lusis AJ, Vulpe CD, Lerch MM, Homuth G, Kacprowski T, Schmidt CO, Nöthlings U, Karlsen TH, Lieb W, Laudes M, Franke A, Huttenhower C. 2019. Obese Individuals with and without Type 2 Diabetes Show Different Gut Microbial Functional Capacity and Composition. Cell Host Microbe 26:252–264.e10.

41. Wang DD, Nguyen LH, Li Y, Yan Y, Ma W, Rinott E, Ivey KL, Shai I, Willett WC, Hu FB, Rimm EB, Stampfer MJ, Chan AT, Huttenhower C. 2021. The gut microbiome modulates the protective association between a Mediterranean diet and cardiometabolic disease risk. Nat Med 27:333–343.

42. Pereira D, Seneviratne CJ, Koga-Ito CY, Samaranayake LP. 2018. Is the oral fungal pathogen Candida albicans a cariogen? Oral Dis 24:518–526.

43. Mayer FL, Wilson D, Hube B. 2013. Candida albicans pathogenicity mechanisms. Virulence 4:119–28.

44. Xiao J, Huang X, Alkhers N, Alzamil H, Alzoubi S, Wu TT, Castillo DA, Campbell F, Davis J, Herzog K, Billings R, Kopycka-Kedzierawski DT, Hajishengallis E, Koo H. 2018. Candida albicans and Early Childhood Caries: A Systematic Review and Meta-Analysis. Caries Res 52:102–112.

45. Falsetta ML, Klein MI, Colonne PM, Scott-Anne K, Gregoire S, Pai CH, Gonzalez-Begne M, Watson G, Krysan DJ, Bowen WH, Koo H. 2014. Symbiotic relationship between Streptococcus mutans and Candida albicans synergizes virulence of plaque biofilms in vivo. Infect Immun 82:1968–81.

46. Falsetta ML, Klein MI, Lemos JA, Silva BB, Agidi S, Scott-Anne KK, Koo H. 2012. Novel antibiofilm chemotherapy targets exopolysaccharide synthesis and stress tolerance in Streptococcus mutans to modulate virulence expression in vivo. Antimicrob Agents Chemother 56:6201–11.

47. Metwalli KH, Khan SA, Krom BP, Jabra-Rizk MA. 2013. Streptococcus mutans, Candida albicans, and the human mouth: a sticky situation. PLoS Pathog 9:e1003616.

48. Ward TL, Dominguez-Bello MG, Heisel T, Al-Ghalith G, Knights D, Gale CA. 2018. Development of the Human Mycobiome over the First Month of Life and across Body Sites. mSystems 3.

49. Diaz PI, Hong BY, Dupuy AK, Strausbaugh LD. 2017. Mining the oral mycobiome: Methods, components, and meaning. Virulence 8:313–323.

50. Dupuy AK, David MS, Li L, Heider TN, Peterson JD, Montano EA, Dongari-Bagtzoglou A, Diaz PI, Strausbaugh LD. 2014. Redefining the human oral mycobiome with improved practices in amplicon-based taxonomy: discovery of Malassezia as a prominent commensal. PLoS One 9:e90899.

51. de Jesus VC, Shikder R, Oryniak D, Mann K, Alamri A, Mittermuller B, Duan K, Hu P, Schroth RJ, Chelikani P. 2020. Sex-Based Diverse Plaque Microbiota in Children with Severe Caries. J Dent Res 99:703–712.

52. Diaz PI, Dongari-Bagtzoglou A. 2021. Critically Appraising the Significance of the Oral Mycobiome. J Dent Res 100:133–140.

53. Frenkel A, Hirsch W. 1961. Spontaneous development of L forms of streptococci requiring secretions of other bacteria or sulphydryl compounds for normal growth. Nature 191:728–30.

54. Kanasi E, Dewhirst FE, Chalmers NI, Kent R, Jr., Moore A, Hughes CV, Pradhan N, Loo CY, Tanner AC. 2010. Clonal analysis of the microbiota of severe early childhood caries. Caries Res 44:485–97.

55. Ling Z, Kong J, Jia P, Wei C, Wang Y, Pan Z, Huang W, Li L, Chen H, Xiang C. 2010. Analysis of oral microbiota in children with dental caries by PCR-DGGE and barcoded pyrosequencing. Microb Ecol 60:677–90.

56. Takahashi N, Yamada T. 1999. Glucose and lactate metabolism by Actinomyces naeslundii. Crit Rev Oral Biol Med 10:487–503.

57. Brailsford SR, Shah B, Simons D, Gilbert S, Clark D, Ines I, Adams SE, Allison C, Beighton D. 2001. The predominant aciduric microflora of root-caries lesions. J Dent Res 80:1828–33.

58. Takenaka S, Edanami N, Komatsu Y, Nagata R, Naksagoon T, Sotozono M, Ida T, Noiri Y. 2021. Periodontal Pathogens Inhabit Root Caries Lesions Extending beyond the Gingival Margin: A Next-Generation Sequencing Analysis. Microorganisms 9.

59. Mishra K, Bukavina L, Ghannoum M. 2021. Symbiosis and Dysbiosis of the Human Mycobiome. Front Microbiol 12:636131.

60. Baker JL, Bor B, Agnello M, Shi W, He X. 2017. Ecology of the Oral Microbiome: Beyond Bacteria. Trends Microbiol 25:362–374.

61. Van Leeuwen M, Rosema N, Versteeg PA, Slot DE, Hennequin-Hoenderdos NL, Van der Weijden GA. 2017. Effectiveness of various interventions on maintenance of gingival health during 1 year - a randomized clinical trial. Int J Dent Hyg 15:e16–e27.

62. Valm AM. 2019. The Structure of Dental Plaque Microbial Communities in the Transition from Health to Dental Caries and Periodontal Disease. J Mol Biol 431:2957–2969.

63. Hwang G, Liu Y, Kim D, Sun V, Aviles-Reyes A, Kajfasz JK, Lemos JA, Koo H. 2016. Simultaneous spatiotemporal mapping of in situ pH and bacterial activity within an intact 3D microcolony structure. Sci Rep 6:32841.

64. Mikx FH, van der Hoeven JS, König KG, Plasschaert AJ, Guggenheim B. 1972. Establishment of defined microbial ecosystems in germ-free rats. I. The effect of the interactions of streptococcus mutans or Streptococcus sanguis with Veillonella alcalescens on plaque formation and caries activity. Caries Res 6:211–23.

65. Delwiche EA, Pestka JJ, Tortorello ML. 1985. The veillonellae: gram-negative cocci with a unique physiology. Annu Rev Microbiol 39:175–93.

66. Do T, Sheehy EC, Mulli T, Hughes F, Beighton D. 2015. Transcriptomic analysis of three Veillonella spp. present in carious dentine and in the saliva of caries-free individuals. Front Cell Infect Microbiol 5:25.

67. Liu G, Wu C, Abrams WR, Li Y. 2020. Structural and Functional Characteristics of the Microbiome in Deep-Dentin Caries. J Dent Res 99:713–720.

68. Munson MA, Banerjee A, Watson TF, Wade WG. 2004. Molecular analysis of the microflora associated with dental caries. J Clin Microbiol 42:3023–9.

69. Aas JA, Griffen AL, Dardis SR, Lee AM, Olsen I, Dewhirst FE, Leys EJ, Paster BJ. 2008. Bacteria of dental caries in primary and permanent teeth in children and young adults. J Clin Microbiol 46:1407–17.

70. Chhour KL, Nadkarni MA, Byun R, Martin FE, Jacques NA, Hunter N. 2005. Molecular analysis of microbial diversity in advanced caries. J Clin Microbiol 43:843–9.

71. Klinke T, Guggenheim B, Klimm W, Thurnheer T. 2011. Dental caries in rats associated with Candida albicans. Caries Res 45:100–6.

72. Fakhruddin KS, Perera Samaranayake L, Egusa H, Chi Ngo H, Panduwawala C, Venkatachalam T, Kumarappan A, Pesee S. 2020. Candida biome of severe early childhood caries (S-ECC) and its cariogenic virulence traits. J Oral Microbiol 12:1724484.

73. Klinke T, Kneist S, de Soet JJ, Kuhlisch E, Mauersberger S, Forster A, Klimm W. 2009. Acid production by oral strains of Candida albicans and lactobacilli. Caries Res 43:83–91.

74. Houck J, Jacobson R, Bass M, Dasilva C, Baumhauer JF. 2020. Improving Interpretation of the Patient-Reported Outcomes Measurement Information System (PROMIS) Physical Function Scale for Specific Tasks in Community-Dwelling Older Adults. J Geriatr Phys Ther 43:142–152.

75. Curto-Manrique J, Malpartida-Carrillo V, Arriola-Guillén LE. 2019. Efficacy of the lift-the-lip technique for dental plaque removal in preschool children. J Indian Soc Pedod Prev Dent 37:162–166.

76. Sridhar S, Suprabha BS, Shenoy R, Suman E, Rao A. 2020. Association of Streptococcus Mutans, Candida Albicans and Oral Health Practices with Activity Status of Caries Lesions Among 5-Year-Old Children with Early Childhood Caries. Oral Health Prev Dent 18:911–919.

77. Marquis RE, Clock SA, Mota-Meira M. 2003. Fluoride and organic weak acids as modulators of microbial physiology. FEMS Microbiol Rev 26:493–510.

78. Anderson M, Grindefjord M, Dahllöf G, Dahlén G, Twetman S. 2016. Oral microflora in preschool children attending a fluoride varnish program: a cross-sectional study. BMC Oral Health 16:130.

79. Marsh PD, Head DA, Devine DA. 2015. Ecological approaches to oral biofilms: control without killing. Caries Res 49 Suppl 1:46–54.

